# The genetic basis of host choice and resting behavior in the major African malaria vector, *Anopheles arabiensis*

**DOI:** 10.1101/044701

**Authors:** Bradley J Main, Yoosook Lee, Heather M Ferguson, Katharina S Kreppel, Anicet Kihonda, Nicodem J Govella, Travis C Collier, Anthony J Cornel, Eleazar Eskin, Eun Yong Kang, Catelyn C Nieman, Allison M Weakley, Gregory C Lanzaro

**Affiliations:** Vector Genetics Laboratory, Department of Pathology, Microbiology and Immunology/University of California, Davis, 95616, USA; Institute of Biodiversity, Animal Health and Comparative Medicine, University of Glasgow, Glasgow, UK; Environmental Health and Ecological Sciences group, Ifakara Health Institute, Ifakara, United Republic of Tanzania; Department of Entomology and Nematology/University of California, Davis, 95616, USA; Department of Computer Science, University of California Los Angeles, 90095 Los 23 Angeles, TX 7871215, USA

**Keywords:** *Anopheles arabiensis*, host-preference, resting behavior, host-preference, association study, population genetics, chromosome inversion

## Abstract

Malaria transmission is dependent on the propensity of Anopheles mosquitoes to bite humans (anthropophily) instead of other dead end hosts. Recent increases in the usage of Long Lasting Insecticide Treated Nets (LLINs) in Africa have been associated with reductions in highly anthropophilic vectors such as *Anopheles gambiae s.s.*,leaving less anthropophilic species such as *Anopheles arabiensis* as the most prominent remaining source of transmission in many settings. *An.arabiensis* is more of a generalist in terms of its host choice and resting behavior, which may be due to phenotypic plasticity and/or segregating allelic variation. To investigate the potential genetic basis of host choice and resting behavior in *An. arabiensis* we performed a genome-wide association study on host choice (human-or cattle-fed) and resting position (collected indoors or outdoors) in the Kilombero Valley, Tanzania. This represents the first genomic/molecular analysis of host choice and resting behavior in a malaria vector. We identified a total of 4,820,851 SNPs, which were used to conduct the first genome-wide estimates of 'SNP heritability' for host choice and resting behavior in this species. A genetic component was detected for host choice (human vs cow fed; permuted *P* = 0.002), but there was no evidence of a genetic component for resting behavior (indoors versus outside; permuted *P* = 0.465). A principal component analysis (PCA) segregated individuals based on genomic variation into three groups which are characterized by differences at the 2Rb and/or 3Ra paracentromeric chromosome inversions. There was a non-random distribution of cattle-fed mosquitoes between the PCA clusters, suggesting that alleles linked to the 2Rb and/or 3Ra inversions may influence host choice. Using a novel inversion genotyping assay, we detected a significant enrichment of the standard arrangement (non-inverted) of 3Ra among cattle-fed mosquitoes (N=129) versus all non-cattle-fed individuals (N=234; १^2^, *p*=0.007). Thus, tracking the frequency of the 3Ra in An. arabiensis populations is important, especially in relation to the emergence of behavioral avoidance(e.g. shifting toward cattle-feeding) in some populations. A better understanding of the genetic basis for host choice in *An. arabiensis* may also open avenues for novel vector control strategies based on introducing genes for zoophily into wild mosquito populations.

**Author summary:** Malaria transmission is driven by the propensity for mosquito vectors to bite people, whilst its control depends on the tendency of mosquitoes to bite and rest in places where they will come into contact with insecticides. In many parts of Africa, *Anopheles arabiensis* is now the only remaining vec 63 tor in areas where coverage with Long Lasting Insecticide Treated Nets is high. We sought to assess the potential for An. *arabiensis* to adapt its behavior to avoid control measures by investigating the genetic basis for its host choice and resting behavior. Blood fed *An. arabiensis* were collected resting indoors and outdoors in the Kilombero Valley, Tanzania. We sequenced a total of 48 genomes representing 4 phenotypes (human or cow fed, resting in or outdoors) and tested for a genetic basis for each phenotype. Genomic analysis followed up by application of a novel molecular karyotyping assay revealed a relationship between *An. arabiensis* that fed on cattle and the standard arrangement of the 3Ra inversion. This indicates that the host choice behavior of *An. arabiensis* has has a substantial genetic component. Validation with controlled host preference assays comparing individuals with the standard and inverted arrangement of 3Ra is still needed.

## Introduction

Blood-feeding insects impose a substantial burden on human and animal health through their role as disease vectors. In particular, mosquito species that feed on human blood pose an enormous public health threat by transmitting numerous pathogens such as dengue virus, Zika virus and malaria, which together kill more than one million people per year [1,2]. Human exposure to pathogens transmitted by mosquito vectors is determined by vector behaviors such as: (1) propensity to feed on humans relative to other animals (anthropophily) and (2) preference for living in close proximity to humans, as reflected by biting and resting inside houses (endophily) [3]. These traits are known to vary within and between the *Anopheles* mosquito species that transmit malaria [3]. It has been demonstrated since the earliest days of malaria transmission modeling [4] that the degree of anthropophily in vector populations is strongly associated with transmission intensity. Furthermore, the extent to which vectors feed and rest inside houses is a critical determinant of the effectiveness of current frontline control strategies including Long-Lasting Insecticide Treated Nets (LLINs) and Indoor Residual Spraying (IRS) which selectively kill mosquitoes that bite and rest indoors [1].

Vector species that are more generalist with respect to host feeding behavior, like*An.arabiensis,* are thought to be better able to persist in areas of high indoor insecticide use. This is because they are more likely to avoid feeding and resting in areas protected by insecticides. For example, several studies in East Africa have shown dramatic declines in the abundance of the highly anthropophilic species *An.gambiae* s.s. relative to *An. arabiensis* in parallel with the use of LLINs [5–13]. Similar changes in vector species composition in response to LLINs have been reported outside of Africa, including in the Solomon Islands where the highly endophagic, anthropophilic *An. punctulatus* has been nearly eliminated by LLINs whereas the more exophilic *An. farauti* remains [14]. Given the importance of mosquito feeding and resting behavior to the effectiveness of disease control and transmission, there is an urgent need to understand the underlying biological determinants of these behaviors and their impact (short and long term) on the effectiveness of the existing frontline interventions.

Environmental heterogeneity has been shown to have a substantial influence on several important vector behaviors [15], including host choice and resting behavior [3]. For example, a recent study in southern Tanzania reported that the proportion of blood meals taken from humans by *An. arabiensis* fell by over 50% when at least one cow is kept at a household [16]. The resting behavior of mosquito vectors in this study was also highly associated with proximity to livestock; the proportion of *An. arabiensis* resting indoors fell by ~50% when cattle were present at the household [16]. Whilst these studies confirm that the environment can influence malaria vector behavior, far less is known about the influence of mosquito genetics on these behavioral phenotypes. An early study by Gillies [17] experimentally investigated the potential heritability of host choice behavior in *An.gambiae* s.l., and showed these vectors significantly increased their preference for cattle hosts (relative to humans) within a few generations of selection. Other early work demonstrated an association between the 3Ra chromosomal inversion and An. Arabiensis found in cattle-sheds [18], and between the 2Rb chromosomal inversion and human-feeding [19]. Understanding the genetic basis for host choice behavior is essential for elucidation of the co-evolutionary forces that stabilize the transmission of vector-borne diseases, and may enable the development of genetic markers that could be used for rapid quantification of the degree of anthropophily in vector populations as required to estimate transmission risk and plan vector control programs [20].

There is evidence from other mosquito taxa that host feeding behavior has a significant genetic component. For example, a recent study linked allelic variation in the odorant receptor gene *Or4* to human-biting preference in the dengue mosquito vector *Aedes aegypti* [21]. However, to date, no ortholog for *AaegOr4* has been identified in Anopheline mosquitoes [22], and no direct functional links between genetic mutations in African malaria vectors and behaviors that influence transmission potential have been identified [3,23–25]. As the genera *Aedes* and *Anopheles* diverged before the emergence of humans (~150MYA) [26], anthropophily likely evolved independently in these species and may involve distinct mechanisms. Developing the ability to track and anticipate shifts in mosquito behaviors such as biting time [27], anthropophily [3], and resting behavior [28] will help improve vector surveillance and anticipation of whether the effectiveness of control measures are being eroded by mosquito behavioral adaptations [29]. Such mosquito behavioral shifts that reduce their contact with interventions, termed behavioral avoidance, may be a significant threat to the long-term goal of malaria elimination [30]. Understanding the genetic contribution to these phenotypes is a critical first step toward effective mosquito control in the future.

Due to the role of *An. arabiensis* in maintaining residual malaria transmission across much of sub-Saharan Africa [8,13,31], we conducted a comprehensive investigation of the genetic basis of host choice and resting habitat choice in this phenotypically variable species. This included the first application of both whole genome sequencing and a population-scale assessment of chromosome inversion frequencies to test for associations between mosquito behavioral phenotypes and genotype. Our aim was to elucidate genetic factors that are associated with mosquito behaviors, and compare potential candidate genes with other important disease vector species such as *Ae. aegypti,* whose preference for humans has been recently described at the genetic level [21]. Additionally, we hope information gathered here can be of use to future malaria control scenarios by highlighting the potential of *An. arabiensis* to evolve behavioral avoidance strategies that could either decrease transmission (e.g. zoophily) or diminish control effectiveness (e.g. outdoor resting).

## Results

### Analysis of host choice

We analyzed the blood meal from 1,731 *An. arabiensis* females that were captured resting indoors or outdoors from 3 villages in Tanzania. Specific hosts were identified using a multiplex genotyping assay performed on DNA extracted from female abdomens (see methods). The relative frequencies of blood meals from a given host varied by site, but cattle was the most abundant blood source detected in all three sites. Lupiro had a significantly higher proportion of human-fed mosquitoes (24%; *P* <0.0001, Fisher exact) than Minepa (7%) and Sagamaganga (11%; Figure 1). Mosquitoes that tested positive for more than one host were rare (<5%; Figure 1). To investigate temporal and spatial variation of host choice, mosquitoes were collected from several households throughout a period of 2 years (Table S9). A backward model selection approach with Generalized Linear Mixed Models (GLMMs) was used to investigate whether host choice was impacted by different environmental factors. Livestock presence at the household level (yes or no),season (dry or wet), village and trapping location (in or out) were included into a maximum model as fixed effects while collection date and household were added as random effects (Table S10). The final model showed that livestock presence at the household level and trapping location (indoor or outdoor) were associated with the frequency of human fed mosquitoes found. The proportion of human fed *An. arabiensis* varied by household and was inversely correlated with the presence of livestock (*P*<0.0001, Coeff=−1.92; GLMM, Table S11). The frequency of human fed mosquitoes was also correlated with trapping location-less human fed mosquitoes were collected in outdoor traps(*P*=0.0083, Coeff=− 0.7349; GLMM, Table S11).

**Figure 1.**
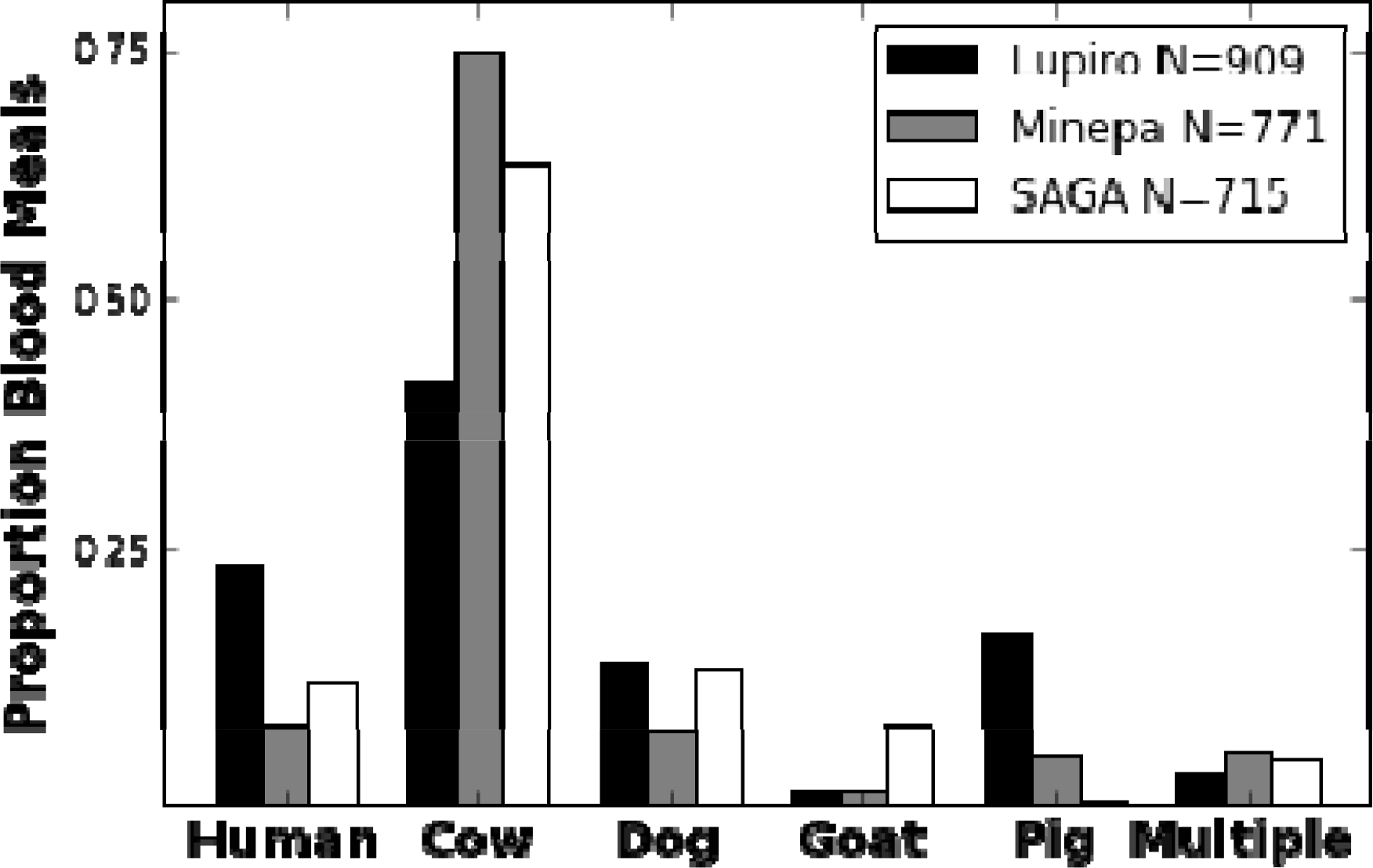
**Relative host choice between villages**. This figure describes the results of bloodmeal analysis of An. Arabiensis collected from: Lupiro, Minepa, and Sagamaganga (SAGA). We detected multiple hosts in <5% of individuals. Different combinations of mixed host bloodmeals were pooled and shown as “Multiple”. A few chicken bloodmeals were also detected in each site (not shown).

### Testing for a genetic component underlying host choice and indoor resting behavior

To elucidate a genetic component to host choice and resting behavior, we sequenced a total of 48 individual *An. arabiensis* genomes (median coverage=18x; Table S2). In terms of host choice, this collection included 25 cattle-fed and 23 human-fed individuals from both indoor (N=24) and outdoor (N=24) resting sites. From these genomes, we identified a set of 4,820,851 segregating SNPs after a minor allele frequency threshold of 10% was imposed. Using these data, we estimated the genetic component (or “SNP heritability” [32]) for each phenotype (see methods). The sample size of 48 genomes was not sufficient to estimate SNP heritability with high confidence. For example, the heritability estimate for human-fed vs. cattle-fed mosquitoes was *H*=0.94, SE=3.47 and indoor vs. outdoor was *H*^2^=0.05, SE=2.34. Thus, we permuted the phenotypes to simulate the null hypothesis of no connection between the SNP data and each behavior. We then compared the estimate of the SNP heritability from the real data with the estimates from each of 10,000 permutations. This test supports the initial heritability estimates indicating a genetic component for host choice (human vs. cow fed; permuted *P*=0.001) and no genetic component for resting behavior (indoor vs. outdoor, permuted *P*=0.470). Due to the lack of evidence for a genetic component for resting behavior, we restricted further analysis to elucidating the observed association between host choice and genotype.

### Genetic structure

To test for the existence of genetic structure within our set of 48 sequenced genomes, individuals were partitioned by genetic relatedness using a principle component analysis on all SNPs (PCA; see methods). Using this approach, we observed 3 discrete genetic clusters (Figure 2a). Genome-wide *F_ST_* in sliding windows between individuals in each PCA cluster (see methods) revealed that the clusters can be explained by distinct combinations of 3Ra and 2Rb chromosome inversion states (Figure 2b). Using a novel inversion genotyping assay (see methods), we determined the 2Rb and 3Ra inversion states for individuals represented in each PCA cluster (2Rb_3Ra): left=bb_a+, middle=bb_++, and right=b+_++. There was an enrichment of cattle-fed mosquitoes among bb_++ individuals (*P*<0.001; Fisher Exact with Freeman-Halton extension).

**Figure 2.**
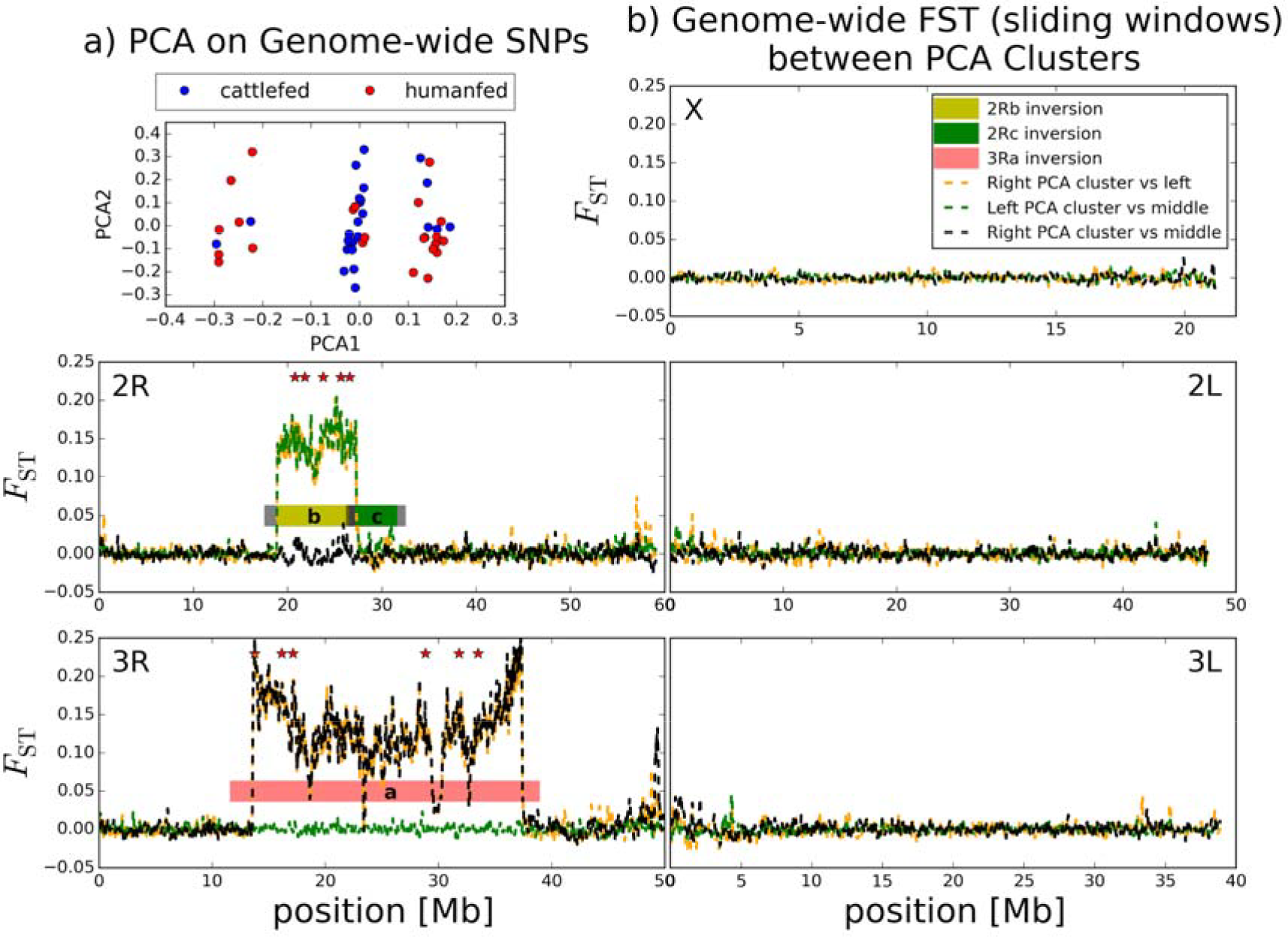
**Genetic variation explained by the 2Rb and 3Ra inversions.** a) Genetic structure was assessed using genome-wide SNP data 579 for individual An. arabiensis females using a PCA analysis. Three discrete PCA clusters were observed. Red = human-fed and blue = cattle-fed. There is an enrichment of cattle-fed individuals in the middle PCA cluster (P < 0.001; Fisher Exact). (b) To reveal differentiated genomic regions underlying the distinct PCA clusters (left, middle, and right) we plotted FST for each chromosome in 100kb windows with 20kb steps between the PCA clusters. The outside PCA clusters differed at the 2Rb and 3Ra inversions (orange), left versus middle PCA clusters differed at 2Rb only (green), and right versus middle differed at 3Ra only (black). Stars indicate the position of SNPs chosen for the inversion genotyping assay.

### Testing for associations between inversion state and host choice

To explore the relationship between the 3Ra and 2Rb inversion state and host choice, we developed and employed a novel inversion genotyping assay. In brief, we selected SNPs near the inversion breakpoints with extreme *F_ST_* values between genomes grouped by distinct 3Ra or 2Rb inversion states. We then genotyped our 11 inversion diagnostic SNPs (3Ra=6, 2Rb=5) in parallel using the Sequenome iPLEX platform (see methods). In total, we genotyped 363 bloodfed females from the village of Lupiro for inversion state. The sample was composed primarily of human-fed (37%) or cattle-fed mosquitoes (36%; Table S7). The 2Rb and 3Ra inversion frequencies were within Hardy-Weinberg (HW) expectations for all samples (*P*=0.55 and 0.90, respectively). However, the 3Ra inversion was outside of HW among dog-fed individuals (*P*=0.02; N=40, Table S7). Only four 3Ra homozygotes were observed (N=363); three fed on dog and one fed on human. The frequency of the 3Ra inversion in Lupiro ranged from 7.94% in cattle to 16.67% in pig-fed mosquitoes. The 2Rb inversion ranged from 81.06% in human to 95% in dog-fed specimens (Table S5). We focused on three major comparisons to test for a relationship between inversion state and host choice: 1)cattle-fed versus human-fed, 2) human-fed versus non-human-fed, and 3) cattle-fed versus non-cattle-fed. After correcting for multiple tests (significant *P*=0.017), there was no evidence for an enrichment of standard arrangement of 3Ra (3R+) in cattle-fed mosquitoes compared to human-fed (*P*=0.047, x^2^; N=263; Table 1) and no relationship was detected between 3Ra and human-fed versus non-human-fed mosquitoes (*P*= 0. 553, x^2^; N=263; Table 1b). However, a significant enrichment of the standard arrangement of 3Ra was observed in cattle-fed versus non-cattle-fed (*P*=0.007, *x*^2^, N=;363; Table 1).

**Table 1.** **Inversion frequencies by host** Mosquitoes were collected from the village of Lupiro. The inversion frequencies (freq a or b) were not calculated for host categories with low sample sizes. The sum of human-and cattle-fed mosquitoes (bottom four categories) included pure (e.g.human) and mixed host (e.g. dog+human) samples.

| Host | ++ | a+ | aa | N | a | + | freq a |
| --- | --- | --- | --- | --- | --- | --- | --- |
| human | 99 | 32 | 1 | 132 | 34 | 230 | 12.88% |
| cattle | 106 | 20 | 0 | 126 | 20 | 232 | 7.9% |
| pig | 38 | 19 | 0 | 57 | 19 | 95 | 16.67% |
| dog | 30 | 7 | 3 | 40 | 13 | 67 | 16.25% |
| goat | 2 | 1 | 0 | 3 | 1 | 5 |  |
| cattle+goat | 2 | 0 | 0 | 2 | 0 | 4 |  |
| human+cattle | 1 | 0 | 0 | 1 | 0 | 2 |  |
| dog+human | 0 | 1 | 0 | 1 | 1 | 1 |  |
| dog+pig | 1 | 0 | 0 | 1 | 0 | 2 |  |
|  |  |  |  | human | 35 | 233 | 13.06% |
|  |  |  |  | non-human | 53 | 405 | 11.57% |
|  |  |  |  | cattle | 20 | 238 | 7.75% |
|  |  |  |  | non-cattle | 68 | 400 | 14.53% |
| Host | ++ | b+ | bb | N | b | + | freq b |
| human | 4 | 42 | 86 | 132 | 214 | 50 | 81.06% |
| cattle | 4 | 33 | 89 | 126 | 211 | 41 | 83.73% |
| pig | 1 | 18 | 38 | 57 | 94 | 20 | 82.46% |
| dog | 0 | 4 | 36 | 40 | 76 | 4 | 95.00% |
| goat | 0 | 1 | 2 | 3 | 5 | 1 |  |
| cattle+goat | 0 | 1 | 1 | 2 | 3 | 1 |  |
| human+cattle | 0 | 0 | 1 | 1 | 2 | 0 |  |
| human+dog | 0 | 1 | 0 | 1 | 1 | 1 |  |
| dog+Pig | 0 | 0 | 1 | 1 | 2 | 0 |  |
|  |  |  |  | human | 217 | 51 | 80.97% |
|  |  |  |  | non-human | 391 | 67 | 85.37% |
|  |  |  |  | cattle | 216 | 42 | 83.72% |
|  |  |  |  | non-cattle | 392 | 76 | 83.76% |

### Candidate genes within 3Ra

Due to the association between host choice and 3Ra, we explored allelic variation in genes in the “odorant binding” gene ontology category (GO:0005549) that occur within the 3Ra breakpoints. To accomplish this,we sorted selected genes by mean *Fst* estimates at each gene level (plus 1kb upstream) between 3Ra standard (N=39) and 3Ra inverted (N=9) genomes (Table S8). Among the genes with the highest *F_ST_* was odorant binding protein antennal(5^th^ highest mean *F_ST_*=0.2) and the odorant receptor *Or65* (10^th^ highest mean *F_ST_*=0.18;Table S8).

## Discussion

In this study, we elucidate the genetic basis of host choice and resting behavior in *An.arabiensis* using whole genome sequencing and a novel chromosomal inversion genotyping assay. We did not detect a genetic component(“SNP heritability”) for resting behavior(endo-versus exo-phily). This could be due to substantial “behavioral plasticity” in this phenotype, which would make this phenotype difficult to detect with field collected samples [33,34], However, a genetic component was detected for host choice based on genome-wide SNP data. Using population-scale inversion genotyping, we show that the 3Ra inversion (or linked alleles) is involved. Identifying functional alleles underlying host choice in *An. arabiensis* is particularly exciting because this species has become the dominant malaria vector in many parts of East Africa, where insecticide use is common [13,35–37]. We highlight two intriguing candidate genes within the 3Ra, including odorant binding protein antennal. The *An. gambiae* ortholog is *Obp5,* which is the highest expressed odorant binding protein (OBP) in female antennae and is significantly overexpressed in female versus male heads [38]. Thus, *Obp5* is likely involved in host seeking behavior, which is female-specific. *Obp5* is also significantly overexpressed in non-bloodfed females compared to 24 hours after blood feeding [39], further implicating its importance in host seeking behavior. We also detected allelic variation in Or65 between 3Ra inversion arrangements. This is a “highly tuned” odorant receptor, that has been shown to be responsive to 2-ethylphenol, a compound found in animal urine [40]. As host choice is directly linked to malaria transmission, elucidating the genetic basis of this behavioral phenotype may lead to innovative tools for vector control. The inversion genotyping assay described herein may be a valuable monitoring tool (e.g. after GMM release or zooprophylaxis), potentially indicating the relative feeding plasticity of a population based on the frequency of 3Ra.

### Associating SNPs with human- and cattle-fed *An. arabiensis*

“SNP heritability” provides an estimate of the correlation between phenotype and genome-wide SNP genotypes from pairs of individuals sampled from a population [32]. A strength of this metric is its robustness to complex phenotypes that are influenced by many small-effect mutations, which may be the case for host choice in *An. arabiensis.*In this study, we collected mosquitoes that were blood-fed and resting indoors or outdoors to assess the genetic basis of host choice and indoor resting behavior. Each phenotype is complex and may be affected, at least in part, by innate preference and the local environment, including host availability and indoor resting sites. Despite our inability to control for environmental heterogeneities in this field experiment, the SNP heritability analysis detected a genetic component for host choice. Due to the low LD (~200bp) across the genome of this species [41], increased samples sizes (e.g. 100-1000) are needed to get a quantitative estimate of the SNP heritability of host choice and potentially uncover additional candidate genes. Larger sample sizes may also uncover a genetic component to resting behavior, which we did not detect here but cannot rule out. Previously, high inversion polymorphism has been detected in *An. arabiensis* in malarious areas in Nigeria with some inversions showing changes in frequencies linked to different geographical areas [42]. This could be linked to selection pressures driven by vector control and/or host availability on resting and feeding behavior.

### Cattle-feeding linked to the 3Ra inversion

A principal component analysis on genome-wide SNPs resulted in 3 discrete clusters distinguishable by the 3Ra and 2Rb inversion (Figure 2). There was no significant enrichment among the 48 sequenced individuals in any given cluster (x^2^; *P*=0.23). However,the distribution of human-and cattle-fed mosquitoes among the clusters was non-random (*P*<0.01; 2x3 Fisher Exact), suggesting that the inversion/s may contain alleles related to host choice. In *An. arabiensis,* indirect associations have also been made between host choice and inversions, like 3Ra in Ethiopia [18] and Kenya [43]. A non-random distribution of the 2Rb inversion has also been reported between human-and cattle-fed mosquitoes [19], but we are unaware of *An. arabiensis* studies with paired karyotype and host choice information from each individual mosquito.

To test for an association between host choice and these inversions with a much larger sample size, we developed a novel inversion genotyping assay (see methods). It should be noted that the inversions represent one or more linked alleles among many possible other contributing alleles throughout the genome. To ensure that our genotyping method was robust, we selected multiple SNPs near the inversion breakpoints for each inversion (see stars in Figure 2b). We associated the inversion genotype results to the standard or inverted arrangement of the 2Rb and 3Ra by genotyping 15 karyotyped samples (Table S5). This allowed us to determine the inversion state and bloodmeal source (host) from each individual in a high-throughput and economical fashion. Further testing is needed to assess how well this assay would perform with *An. arabiensis* samples from outside our study sites in Tanzania.

Using this molecular karyotyping method, we observed an enrichment of the standard arrangement of 3Ra among cattle-fed mosquitoes (*p*=0.007, x^2^, N=363; Table 1b). The frequency of the 3Ra inversion in dog-fed, goat-fed, and human-fed mosquitoes was substantially higher than cattle-fed mosquitoes (Table 1). Overall, the frequencies of 3Ra were within Hardy Weinberg expectations (*P*=0.55; Table S7). Notably, there was an enrichment of 3Ra/a homozygotes among dog-fed mosquitoes (*P*=0.02; Table S7). These genotypes are so rare in Tanzania, some have postulated the presence of a recessive lethal in 3Ra [44]. The enrichment of 3R+ among cattle-fed mosquitoes is strong support for a genetic component to host choice, which is consistent with the report that zoophily can be selected for [17]. The fact that all other species in the *Anopheles gambiae* species complex are fixed for the standard arrangement of 3Ra, strongly suggests that 3Ra is derived [45]. Thus, one possible explanation for this pattern in *An. arabiensis* is that the standard arrangement of 3Ra (3R+) is the ancestral state and alleles therein facilitate specialization on cattle. A loss-of-function mutation was then acquired early on in a gene or gene network that is critical to specializing on bovids in the haplotype representing the inverted arrangement of 3Ra. As a result, individuals with the 3Ra are more opportunistic feeders. This hypothesis is consistent with behavioral heterogeneities and 3Ra frequencies across Africa. For example, *An. arabiensis* is reportedly more anthropophilic in West African countries like Burkina Faso and Mali [46,47], where the frequency of 3Ra is very high (~40–60%; [48–51]) compared to East African populations, like our field site in Tanzania (~12% or less), and others [18,43,52–54]. The diversity of host feeding behaviors, including both generalists (e.g. *An. arabiensis* with 3Ra) and specialists (e.g. *An. gambaie s.s.)* among species in the *An. gambiae* complex make this a fascinating system to study the evolution of host preference.

While we provide strong evidence for a role of allelic variation within 3Ra underlying *An. arabiensis* host choice, the effect size (i.e. relative contribution to the phenotype) is unclear. Correcting for environmental variation is likely very important when choosing representative samples for each genotype. For example, a human-fed mosquito may be more meaningful if there is an abundance of alternative hosts nearby (e.g. cattle). This was shown by Tirados et al. [55], where *An. arabiensis* was found to persistently bite humans despite being surrounded by cattle, negating a zooprophylactic effect of cattle. This highlights the importance of integrating genetic analyses into a wider context. Colony-based host preference assays involving representatives from each 3Ra inversion state in a controlled environment may be the most effective way forward. Previous tests for population structure only revealed differentiation between distant villages [41]. Thus, by comparing individual genomes representing host choice phenotypes (and resting behavior) from within the same village (Lupiro), we limited the identification of demographic SNPs in our data set. However, to assess the role of 3Ra more broadly, additional studies involving study sites across the range of *An. arabiensis* are needed.

### Environmental component to host choice

In this study, mosquitoes were sampled directly from the field, which enabled us to examine the contribution of environmental heterogeneity to the host choice phenotype.
Cattle was the preferred host at each collection site, but we found differences in relative host-choice patterns between villages. For example, the human-fed mosquitoes were significantly more common in Lupiro (24%) versus in Minepa (7%) and Sagamaganga (11%, Fisher exact P<0.0001). This trend varied by collection year (Table S2). The decreased human blood index in households with livestock nearby (Table S11) and lower frequency of human-fed mosquitoes collected in outdoor traps, where both cattle and humans are available, indicates that local host availability has a major influence on host choice and is consistent with previous reports [16]. As individuals with the standard arrangement of 3Ra appear to prefer cattle (Table 1), the effect of host availability on host choice will likely be stronger in populations where the inverted arrangement of 3Ra is relatively rare.

### Future directions

This study presents important data suggesting a genetic component to host choice in the malaria vector *An. arabiensis.* We show that the 3Ra inversion is involved, at least in part. This association and the introduction of a novel inversion genotyping assay may be a valuable tool for future malaria control strategies involving *An. arabiensis.* For example, tracking the frequency of the 3Ra in *An. arabiensis* may elucidate the emergence of behavioral avoidance (e.g. shifting toward zoophily) so countermeasures can be implemented. A better understanding of the genetic basis for host choice in *An. arabiensis* may also improve vector control if cattle-biting mosquitoes can be genetically engineered and released in the population, having an effect similar in concept to zooprophylaxis [56].

Due to the existence of substantial genetic and environmental components to the host choice phenotype in *An. arabiensis,* an important next step from this study would be to establish *An. arabiensis* colonies, ideally from Lupiro, that are representative of each inversion state and perform choice assays in controlled environmental conditions.

## Materials and Methods

### Mosquito collection area

The mosquitoes were collected within 3 villages in the Kilombero River Valley in southeastern Tanzania: Lupiro (S08°23.2956′; E036°40.6122′), Minepa (S08°16.4974′; E036°40.7640′) and Sagamaganga (S08°03.8392′; E036°47.7709′). The Kilombero Valley is dominated by irrigated and rain-fed rice paddies and maize fields bordered by woodland. The annual rainfall is 1200−1800 mm with two rainy seasons. The average daily temperatures range between 20°C and 33°C. Most people in this area are subsistence farmers and/or livestock keepers. Mud or brick houses stand in clusters among a few trees or banana trees. If a household owns livestock, the animals are kept outside a few meters away from the house in sheds (pigs and goats) or within simple cattle fences. Animal sheds with walls and a roof were considered indoor resting areas. Inside houses you will regularly find chickens, cats and sometimes dogs. The mosquitoes will encounter bed nets inside almost all houses in the valley, but no repellents are currently used by people outdoors [57] and livestock are not treated with insecticide [58]. Malaria is endemic in these communities and although prevalence is declining, almost all inhabitants have antibodies for the disease [59]. The dominant malaria vector species are *An. arabiensis* and the *An. funestus* group [60].

### Collection methods

In each village, households chosen for collection were within 100−200m of one another. Indoor mosquito collection method was aspiration using a standard battery-powered CDC Back Pack aspirator (BP, Model 1412, John Hock, Florida USA) [61]. In these collections, the aspirator was used to collect mosquitoes from the main bedroom by sweeping the nozzle over the interior walls, roof and furniture for a fixed period of ten minutes. BP collections were timed to standardize sampling effort across houses. A resting bucket trap (RBu) was used to trap mosquitoes outdoors. The RBu is made from a standard 20 liter plastic bucket lined with black cotton cloth, and set by placing it on its side with the open end facing a house at a distance of approximately 5m. A small wet cloth is placed inside the bucket to increase humidity. Mosquitoes resting inside RBus were collected at dawn by placing the nozzle of a battery-powered modified CDC backpack aspirator at the open end of the bucket and aspirating for 10−20 seconds.

### Ethics

Before collection, meetings were held with community leaders in all villages during which they were informed about the purpose of the study and their participation requested. After their permission had been granted, the study team visited each village and informed consent was obtained from each head of household where trapping was conducted. Research clearance was obtained from the institutional review board of Ifakara Health Institute in Tanzania (IHI/IRB/No: 16−2013) and by the National Institute for Medical Research in Tanzania (NIMR/HQ/R.8c/Vol. II/304).

### DNA extraction

For each specimen, the abdomen was separated from the head and thorax; DNA was extracted separately from each using the QIAGEN Biosprint 96 system and QIAGEN blood and tissue kits (QIAGEN, Valencia, CA). *Anopheles arabiensis* samples were distinguished from other *An. gambiae* s.l. species complex members with the Scott polymerase chain reaction assay [62] and their DNA content was quantified using the Qubit 2.0 Fluorometer (Life technologies, Grand Island, NY).

### Bloodmeal analysis

The specific host species that each mosquito had fed upon was determined by a multiplex genotyping assay on DNA extracted from abdomens [63]. This multiplex genotyping assay can distinguish between blood from cattle, goat, pig, dog, chicken and human.

### Analysis of host choice

Statistical analysis was conducted to compare the proportion of human-fed mosquitoes in total between villages and of these the proportion caught resting indoors using the statistical software R (Core-Team RD, 2013). Variation in the proportion of human-fed *An. arabiensis* within the total catch was investigated. Samples found to contain any human blood represented one category and those containing animal blood another. Generalized linear mixed effects models (GLMM, package lme4 in R [64]) were used, with human-fed mosquitoes versus animal-fed mosquitoes as a response variable with a binomial distribution and fitting village and livestock presence as fixed effects, and date and house of collection as random effects. To be able to explore the resting behavior of *An. arabiensis,* only mosquitoes resting in houses or outdoors but not those caught resting in animal sheds were used for analysis. Here the GLMM were fitted for each village separately with human-fed mosquitoes caught indoors versus outdoors as a response variable with a binomial distribution and livestock as fixed effect and date and house of collection as random effects.

### Cytogenetic analysis

To identify 3Ra, 2Rb, and 2Rc chromosomal inversions, polytene chromosomes were extracted from ovarian nurse cells from half gravid indoor resting mosquitoes using the protocol described by Hunt [65]. Chromosome banding patterns were examined using a Nikon Eclipse e600 phase contrast microscope. The genotypes of the chromosome inversions were scored for each individual mosquito. Photographic images of chromosomes for the majority of individual mosquitoes used in this study are available on PopI OpenProject page - AaGenome
(https://popi.ucdavis.edu/PopulationData/OpenProjects/AaGenome).

### Genomic library preparation and sequencing

To avoid identifying SNPs associated with demography or other environmental factors, we chose to sequence mosquitoes collected from only one village, Lupiro. We focused on this village because it had sufficient human-fed mosquitoes for testing (Figure 1). Genomic DNA was quantified using a Qubit 2.0 fluorometer (Life Technologies). We used 25−50ng of input DNA for library construction. DNA was then cleaned and concentrated with the DNA Clean and Concentrator kit (Zymo Research Corporation). Library preparations were made with the Nextera DNA Sample Preparation Kit (Illumina), using TruSeq dual indexing barcodes (Illumina). Libraries were size-selected with Agencourt AMPure XP beads (Beckman Coulter). We assessed the insert size distribution of the final libraries using a QIAxcel instrument (Qiagen, Valencia, CA) or Bioanalyzer 2100 (Agilent), and the final library concentration was measured with a Qubit 2.0 fluorometer (Life Technologies). Individually barcoded libraries were sequenced with the Illumina HiSeq2500 platform with paired-end 100 base pair reads, at the QB3 Vincent J Coates Genomics Sequencing Laboratory at UC Berkeley. See Table S1 for raw sequence output per sample.

### Genome sequence mapping and SNP identification

We assessed the quality of our genome sequencing reads using the FastQC software (http://www.bioinformatics.babraham.ac.uk/projects/fastqc/). Adaptor sequences and poor quality sequence were trimmed from the raw Illumina Fastq reads using the Trimmomatic software, version 0.30 [66], with default options. Reads were aligned with BWA-mem [67] to the assembled *An. arabiensis* reference genome version AaraCHR (generously provided by Xiaofang Jiang, Brantley Hall, and Igor Sharakhov. Also see [68]). We used the MarkDuplicates module from Picard tools to remove PCR duplicates and the Genome Analysis Tool Kit (GATK) v1.7 to realign reads around indels [69]. The resulting sorted BAM (Binary sequence Alignment/Map) files containing sequences for each read and its mapping position were then used to make a VCF (Variant Call Format) file using samtools (v1.1−12) 'mpileup' and bcftools (v1.1−36) multiallelic-caller. We removed indels using VCFtools (v0.1.13; “−−remove-indels”) and filtered for variable sites using a minor allele frequency threshold of 0.10 (“−−maf 0.1”) and a major allele threshold of 0.9 (“−−max-maf 0.9”).

### Estimating SNP heritability of each phenotype

Host choice and resting behavior phenotypes may be influenced by many small-effect mutations across the genome. SNP heritability is the correlation between the genome-wide genotypic variation and phenotypic variance (V(G)/V(p)). To estimate SNP heritability, the VCF file containing genome-wide SNP data for all samples was converted to PLINK with VCFtools (command “vcftools−−plink”) and then binary ped files (GCTA option: “−−make-bed”) for analysis with the Genome-Wide Complex Trait Analysis software (GCTA; [70]. To calculate “SNP heritability” with GCTA, we first generated a genetic relationship matrix. Then we calculated SNP heritability for host choice (estimated human-fed prevalence=20%) and resting behavior (estimated indoor prevalence=43%). To estimate the permuted p−value, we used a custom python script to randomly permute the phenotype key for 10000 iterations (see supporting information). The permuted p−value was estimated from the proportion of heritability estimates from the randomly permuted phenotype key that were greater than the heritability estimate from the real data.

### Chromosomal inversion genotyping assay

We used GCTA [70] to perform a principal component analysis (PCA) on all whole genome sequenced individuals from Lupiro. This partitioned the individuals into at least three clusters. Genomic differentiation among the three clusters was concentrated in regions corresponding to 2Rb and 3Ra inversions (Figure 2). We identified candidate diagnostic SNPs between the three clusters using F_ST_ values. We selected 6 diagnostic SNPs for 3Ra that span 19.76Mbp, and 5 diagnostic SNPs for 2Rb spanning 6Mbp (Table S4). A multiplex SNP genotyping assay was designed for an iPLEX assay platform using Sequenom Typer AssayDesigner program (Sequenom, Table S3). The Veterinary Genetics Laboratory at UC Davis performed Genotyping using the Sequenom iPLEX.

### Data accessibility

The genetic information and meta data associated with this study are available on dryad and on the open source online vector database PopI: AaGenome (https://popi.ucdavis.edu/PopulationData/OpenProjects/AaGenome/).

## Competing interests

The authors declare that they have no competing interests.

## Author's Contributions

BJM conducted the experiment, data analysis and wrote manuscript. YL and GCL conceived the experiment, conducted field collections, and wrote the manuscript. HF conceived the overall study and helped with the manuscript. KSK coordinated and conducted field collections, analyzed data and contributed to the manuscript. TCC and EYK conducted data analysis. EE performed data analysis and helped write the manuscript. AJC conducted field collections and cytogenetic analysis. AK helped coordinate and implement field studies and mosquito databas 556 es. NJG contributed to the manuscript. CCN and AMW performed molecular work.

## Acknowledgments

We thank Julia Malvick at the Veterinary Genetics Laboratory for her assistance in iPLEX SNP genotyping and three anonymous reviewers for their comments on the previous version of this manuscript. We also thank the inhabitants of Lupiro, Minepa and Sagamaganga for their collaboration during field sampling. Sequencing was performed by the Vincent J. Coates Genomics Sequencing Laboratory at UC Berkeley, supported by NIH S10 Instrumentation Grants S10RR029668 and S10RR027303. Financial support was provided, in part, by the National Institutes of Health grants R01AI085175-03 and T32AI074550.

